# A Web/Cloud based Digital Pathology Platform Framework for AI Development and Deployment

**DOI:** 10.1101/2022.11.04.514741

**Authors:** Zeynettin Akkus, Bryan Dangott, Aziza Nassar

## Abstract

Digitization of glass slides has brought several opportunities with it for computational pathology and artificial intelligence (AI). The application of AI in digital pathology slides shows potential for QA/QC, triaging cases, and assisting pathologists in clinical decision making. We present an extensible and modular web/cloud based digital pathology framework for AI development and deployment. The proposed platform supports collaborative multi-user and multi-device annotation, remote slide access, and remote telepathology or teleconsultation tasks.

## INTRODUCTION

Digital pathology is the process of digitizing a specimen glass slide with a whole slide scanner, creating an image file with a pyramidal scheme of multiple resolutions, managing, sharing of digital slides, and viewing and reading cases though computer monitors and mobile devices^1,2^. Digital pathology has potential to improve patient care by enabling rapid referral of cases and sharing for expert opinions across pathology networks and organizations, increase efficiency of the lab workflow promote education and training of lab specialist, and create vast number of opportunities for artificial intelligence (AI) to optimize, advance, automatize pathology services. However, a significant amount of investment is necessary for transforming pathology services with digital pathology such as supporting IT infrastructure, staffing, training, and AI computing resources and integration^3–8^.

We propose an extensible and modular digital pathology platform framework that allows flexible and scalable AI development and deployment and its integration to the research and clinical environment.

## METHODS

We built a web/cloud based digital pathology platform that allows slide storage and management in local and remote sites, slide viewing and annotation through a web viewer ^9,10^, and development and deployment of AI models through google cloud computing services^11^. The framework of our platform includes several components such as database for content management, front-end web viewer and annotation, back-end APIs for streaming slides and AI model results to client-side.

### The FRAMEWORK OF PLATFORM

- Whole Slide Image (WSI) Viewer and Annotation tools
- Support All Proprietary WSI and DICOM WSI Images ^11–14^
- AI model Development and Deployment pipelines through Google Cloud
- Integrated to Sectra Medical PACS^15^
- Remote File Access and Distributed Service Model
- Integrated Manual/Automated Scanner^16^
- Rapid Onsite Evaluation using AI and Telepathology/Teleconsultation
- Integrated to Cloud DICOM Stores through Google Healthcare API^15,17^

**Figure 1:**
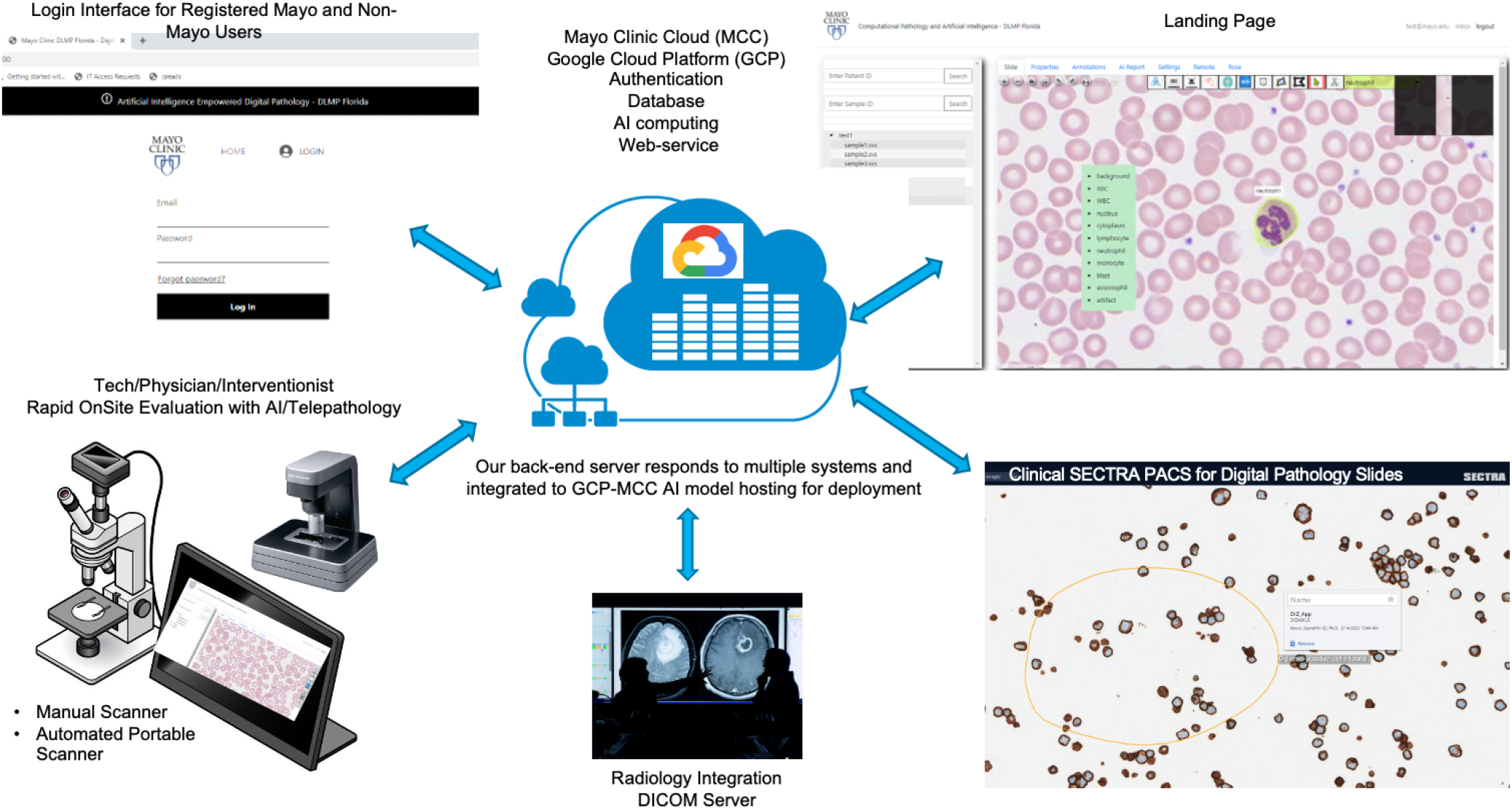
A Practical Digital Pathology Framework for AI Development and Deployment

## APPLICATIONS AND RESULTS

Several AI applications have been developed with this platform. We are going to present two use cases:

1. Rapid On-Site Evaluation of Endobronchial Ultrasound Guided Aspirations using AI. 2) Objective Quality Control of Digital Pathology Slides using AI.

### 1) Rapid On-Site Evaluation of Endobronchial Ultrasound Guided Aspirations using AI

We have developed an artificial intelligence (AI) solution for ROSE of digital pathology slides of EBUS specimens for assessment of sample adequacy and preliminary diagnosis. We designed a custom AI model based on a convolutional neural network that detects tumor cell clusters and is trained to predict adequacy on lymph nodes and subtypes of cancer. Our study population includes 33 digital pathology slides of lung lymph node biopsy from 33 patients with the distribution of 6 adenocarcinoma, 4 lymphoma, 3 neuroendocrine carcinoma, 3 small cell carcinoma, 4 squamous cell carcinoma, 4 other cancer types, 3 non-diagnostics and 6 negative cases. We randomly selected hot areas in digital slides and annotated diagnosable tissue regions. We split our data into 80% for training and 20% for validation of our model.

Nearly 1000 annotation tiles were created. The training performance of our model was 96.6% accuracy level for detecting clusters of tumor cells and excluding anything else. The performance of the model predicting cancer subtypes was 95.3% accurate. Our model performance on validation set (95.7% accuracy) was in concordance with the training performance in tumor cell detection but lower in cancer subtype prediction (55.8% vs. 95.3% accuracy). An example is shown in Figure 2.

**Figure 1a:**
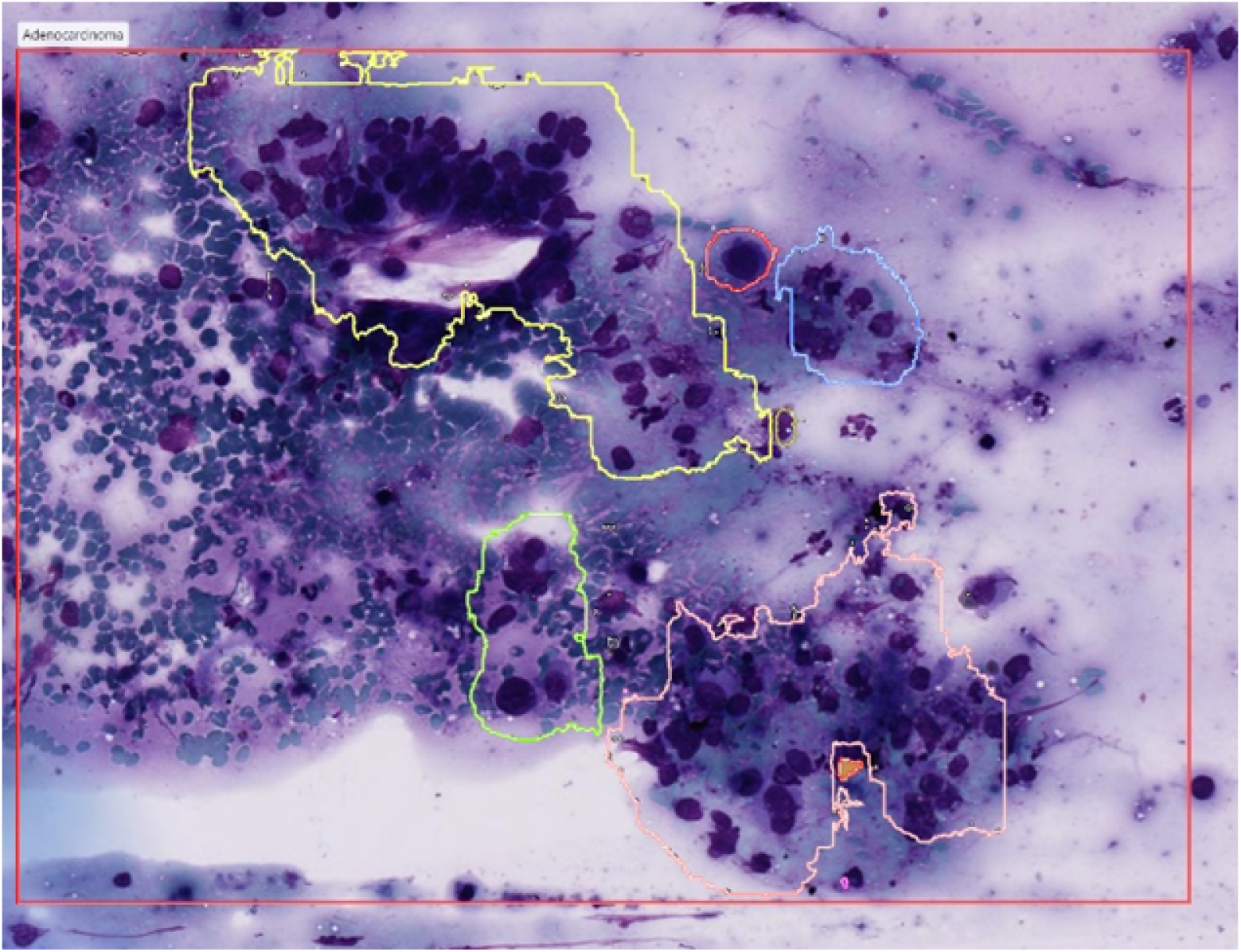
An example of model predictions for tumor cell detection and type prediction.

### 2) Objective Quality Control of Digital Pathology Slides using AI

In this study, we experimented with several well-known convolutional neural networks (CNN) (i.e., resnet50, inceptionv3, inception_resnet, densenet, and VGG CNN architectures) to predict the presence of artifacts in whole slide image tiles. We performed a grid search with a range of values to find the optimal tile size (e.g., 128×128, 256×256, and 512×512 pixels) and optimal magnification level (e.g., from 2.5x to 20x) to detect specific artifacts. We collected a total of 78 whole slide images and annotated the artifacts present in each slide. The data was split into a 7:1:2 ratio in slide level for training (56 slides), test (9 slides), and validation sets (13 slides), respectively. We trained CNN models to predict 14 classes: background (normal), dust, focus, pen, coverslip edge, fold, chatter, dropped tissue, air bubble, tissue scratch, coverslip scratch, debris, and others (figure 3).

**Figure 3.**
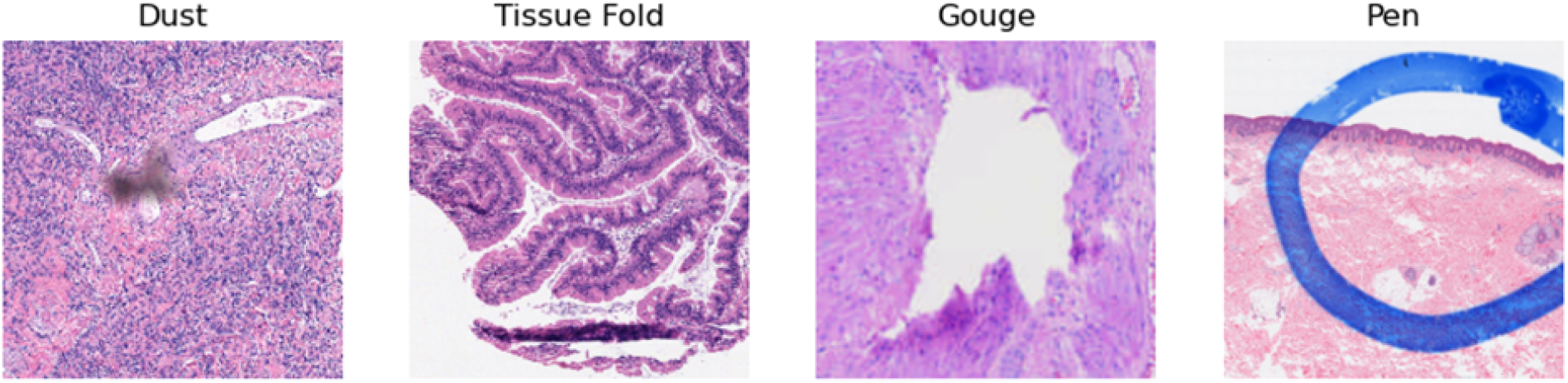
Example artifacts

The tile size of 256×256 pixels gave the overall best performance while a larger tile size gave better performance for the dropped tissue category. Low magnification was often satisfactory for artifacts such as dust, pen, and tissue folds but a higher magnification is necessary to detect out-of-focus artifacts. Inception_resnet was the best performing model with overall accuracy of 66% and 65% for test and validation sets, respectively. The performance of the model jumps to 74% and 80% accuracy if true prediction is among top two predictions of the model.

## DISCUSSION

- Our cloud-based platform allows instant switching between AI models for application to any WSI.
- Interactive predictions in real time are achievable in selected regions of a WSI.
- The processing time of WSI is dependent on the size of the input image tile, magnification level, and the parallelization configuration of Graphics processing units (GPUs).
- Distributed system model allows accessing any slide at any site with a little effort.
- GCP provides load balancing and scalability to handle multiple user access and AI jobs.
- Extensibility and modularity are key to provide tailored solutions to pathology labs
- Creating secure API connections is an important component of handling sensitive healthcare data and services. Google cloud Apigee^18^ is used as an API gateway to provide abstraction and security for backend API services.
- Sharing and streaming WSI in a de-identified manner is important for accessing multi-center data and developing generalizable AI models. Our platform could also be used for federated learning approaches across multi-national and multi-institutional datasets.
- AI-empowered digital pathology will promote education, allow training lab specialists in a shorter time, and serve as a content management and generator service.

## CONCLUSION

- Our platform enables pathologists by supporting collaborative, multi-user, and multi-device annotations for AI model development.
- The architecture can host a variety of AI models and can be deployed to access slides distributed globally in a de-identified manner.
- Algorithms run on cloud infrastructure can be run prior to slide viewing or interactively with user input.
- The modular and distributed framework model allows flexibility to add or remove modules to accommodate future growth.

